# Site-specific PEGylation crosslinking of L-asparaginase subunits to improve its therapeutic efficiency

**DOI:** 10.1101/317040

**Authors:** Josell Ramirez-Paz, Manoj Saxena, Louis J. Delinois, Freisa M. Joaquín-Ovalle, Shiru Lin, Zhongfang Chen, Virginia A. Rojas-Nieves, Kai Griebenow

## Abstract

L-Asparaginase is an enzyme successfully being used in the treatment of acute lymphoblastic leukemia, acute myeloid leukemia, and non-Hodgkin’s lymphoma. However, some disadvantages still limit its full application potential, e.g., allergic reactions, pancreatitis, and blood clotting impairment. Therefore, much effort has been directed at improving its performance. A popular strategy is to randomly conjugate L-asparaginase with mono-methoxy polyethylene glycol, which became a commercial FDA approved formulation widely used in recent years. To improve this formulation by PEGylation, herein we performed cysteine-directed site-specific conjugation of the four L-asparaginase subunits to prevent dissociation-induced loss of activity. The conjugation sites were selected at surface-exposed positions on the protein to avoid affecting the catalytic activity. Three conjugates were obtained using different linear PEGs of 1000, 2000, and 5000 g/mol, with physical properties ranging from a semi-solid gel to a fully soluble state. The soluble-conjugate exhibited higher catalytic activity than the non-conjugated mutant, and the same activity than the native enzyme. Site-specific crosslinking of the L-asparaginase subunits produced a higher molecular weight conjugate compared to the native tetrameric enzyme. This strategy might improve L-asparaginase efficiency for leukemia treatment by reducing glomerular filtration due to the increase in hydrodynamic size thus extending half-live, while at the same time retaining full catalytic activity.

## Introduction

L-Asparaginase is a homo-tetramer enzyme that catalyzes the hydrolysis of asparagine to aspartic acid and ammonia [1]. Some types of blood cancers, e.g., acute lymphoblastic leukemia and non-Hodgkin lymphoma need to acquire L-Asn from extracellular sources due to a deficiency in express asparagine synthetase. L-asparaginase has been used as the main therapeutic agent for more than 40 years to treat leukemia and lymphoma [2]. Once injected into the bloodstream L-asparaginase maintains the pool of circulating asparagine under a concentration that affects protein biosynthesis in the malignant cells, without affecting the normal cells, to eventually promote apoptosis [2-6]. In addition to its medicinal application, L-asparaginase has been recently studied as a possible solution to reduce the content of acrylamide in cooked food products. Acrylamide was cataloged as “probably carcinogenic to humans” by the International Agency for Research on Cancer, besides being a toxin. Treating raw products with L-asparaginase before cooking reduces the main acrylamide precursor asparagine, which reduces acrylamide content in cooked products [7].

The main limitation during medical treatments with L-asparaginase is its inactivation due to degradation by proteolytic enzymes and anti-asparaginase antibodies developed by the host, which cause rapid serum clearance of the drug [2-6,8,9]. A common strategy to improve the pharmacokinetics of protein-based drugs is their modification with polyethylene glycol. PEGylation extents the half-life of proteins by increasing their molecular weight to retard glomerular filtration, while simultaneously hiding antigenic and proteolytic epitopes through its known stealth effect [10-12]. A commercial formulation of *E. coli* L-asparaginase II modified with mono-methoxy polyethylene glycol exhibited about five-fold prolonged blood half-life compared to the non-modified enzyme in clinical studies, and a significant reduction in the incidence of neutralizing antibodies was also observed [3-6,13]. Such improvements reduce the frequency of injections and dosage during treatment [5,6].

PEGylation is an efficient method to improve protein pharmacokinetics but is usually accompanied by adverse effects on the pharmacodynamics [10-12], e.g., an increasing degree of PEGylation is correlated to the reduction of catalytic activity and substrate affinity due to restriction of protein structural dynamics and steric effects [14,15]. In the case of L-asparaginase catalytic activity was drastically reduced to 8% compared to the natural enzyme when around half of the lysine residues were PEGylated [16-22]. Furthermore, a clinical study indirectly revealed that the *in vivo* substrate affinity was affected, since higher concentration of PEGylated L-asparaginase as compared to the non-modified enzyme was needed to maintain the serum concentration of L-Asn under ~10 μM [13], which is relevant since the *in vivo* substrate affinity of this enzyme is 29 μM and the normal steady-state asparagine concentration in serum is around 60 μM [4,23]. Another disadvantage is the discovery of anti-PEG antibodies due to the high exposure to this polymer that we experience in daily life [24-26].

While it is indisputably necessary to investigate other types of polymers and methods to improve the pharmacodynamics of protein-based drugs, over two decades of clinical use has stablished PEGylation as the most efficient method to extent biopharmaceutics half-life, besides being an affordable technology still evolving [27]. As for the future of L-asparaginase, new formulations are constantly entering clinical trials [2,3], and there is convincing evidence of its efficiency for more than 40 years [28,29]. For example, a recombinant form from *E. coli* recently entered the European market [2]. An attractive way to modify L-asparaginase pharmacokinetics while at the same time maintaining its pharmacodynamics is by site-specific PEGylation, that is the attachment of PEG polymers at pre-selected sites on the protein surface [30-33]. By attaching PEGs at specific positions, one can avoid obstructing the active site and limit the degree of PEGylation to maintain protein structural dynamics thus to retain full catalytic activity [14,15]. This strategy has been partially validated for L-asparaginase through PEGylation at its natural disulfide bond without losing catalytic activity, independently of the PEGs length [34,35].

In this work we report for the first time the simultaneous site-specific PEGylation and intramolecular crosslinking of L-asparaginase subunits at pre-selected canonical cysteines introduced by mutagenesis. The advantage of this approach is that not only the degree of modification is kept to a minimum in order to retain catalytic activity, but careful selection of the PEGylation positions also offers the possibility to target potential proteolytic and immunogenic epitopes. Moreover, the molecular weight was increased at least four-fold compared to the native L-asparaginase with as low as one PEG molecule per protein subunit and this significantly reduces the chance for glomerular filtration and anti-PEG binding. PEGylation of this therapeutically important enzyme at canonical amino acids is not reported, even though the conjugation chemistry at cysteine residues is well known and highly specific [36]. The most convenient way to express recombinant L-asparaginase is by secretion into the periplasmic space or culture medium [37-40], since cytoplasmic expression leads to formation of inclusion bodies making the purification tedious [41]. However, expression of canonical cysteines affects L-asparaginase secretory expression [42]. Herein we were able to express a double-Cys mutation of L-asparaginase as a secreted product, to later perform the site-specific crosslinking of the subunits. Our findings will benefit the evolving technological improvement of L-asparaginase as therapeutic agent by setting the prove-of-concept of this alternative PEGylation strategy.

## Materials and methods

### Materials

The plasmid pET-22b(+) was purchased from Novagen (Darmstadt, Germany). The vector pET22b-AspII was synthetized by GenScript (Piscataway, NJ, USA). Gibson’s assembling kit was purchased from New England BioLabs (Ipswich, MA, USA). Primers were synthetized and DNA sequencing was performed by MCLAB (San Francisco, CA, USA). The *E. coli* BL21(DE3) competent cells, yeast extract, tryptone, glycerol, tris-base, hydrochloric acid, ampicillin, isopropyl β-D-1-thiogalactopyranoside, sucrose, ethylenediaminetetraacetic acid, magnesium sulfate, vacuum filters, ammonium sulfate, sodium chloride, spin ultrafiltration filters, potassium phosphate monobasic and dibasic, tris-2-carboxyethyl-phosphine hydrochloric, 5,5’-dithiobis-2-nitrobenzoic acid, dithiothreitol, ethanol absolute, acetic acid glacial, Nessler’s reagent, trichloroacetic acid, L-asparagine, natural and randomly-PEGylated L-asparaginase, and Corning clear-bottom 96-well plates, were purchased from Millipore Sigma (St. Louis, MO, USA). The MonoQ, Superdex 200 10/300 GL, and Sephadex G-25 columns were from GE Healthcare Bio-Sciences (Pittsburgh, PA, USA). The MS standards Cal Mix 3 and Glu-Fib-B1 were purchased from SCIEX (Redwood City, CA, USA). The Bi-MAL-PEG linkers were purchased from Creative PEGWorks (Chapel Hill, NC, USA). The 4-20% gradient polyacrylamide gels were purchased from Bio-Rad (Hercules, CA, USA). The Coomassie Brilliant Blue G-250 dye, and Pierce^TM^ BCA protein assay kit were purchased from Thermo Fisher Scientific (Grand Island, NY, USA). Gel densitometry analysis was performed with Gel Analyzer II by Dr. Istvan Lazar (Copyright 2010).

### Cloning

The ansB gene encoding the mature *E. coli* L-asparaginase II along with its natural signaling peptide [43], was synthetized and cloned into the plasmid pET-22b(+) at Nde-I and BamH-I restriction sites to generate the expression vector pET22b-AspII. The L-asparaginase mutants C77S-C105S and A38C-T263C were generated using the Gibson’s assembling method [44]. Commercial DNA sequencing was used to verify the constructs.

### Secretory expression

*E. coli* BL21(DE3) competent cells were transformed with pET22b-AspII and grown in terrific broth medium (24 g/l yeast extract, 12 g/l tryptone, 4 ml/l glycerol, 100 mM potassium phosphate buffer pH 7.2, 100 μg/ml of ampicillin), which is suitable for asparaginase extracellular secretion [37]. Cultures were grown to an OD_600nm_ ~0.200 and tested for induction with 10, 100, 500, and 1000 μM of isopropyl β-D-1- thiogalactopyranoside (IPTG) harvested 4 h post-induction. Secretion of asparaginase catalytic activity was also followed at 8, 16, and 24 h post-induction.

Secretion into the periplasmic space and culture medium was assayed by measuring the presence of asparaginase catalytic activity in the extracytoplasmic compartments. The culture medium fraction was defined as the total activity found in the clear culture medium supernatant after centrifugation. The periplasmic space fraction was obtained by osmotic shock as follows. The cell pellet was suspended in osmotic solution 1 (20 %w/v sucrose, 1 mM EDTA, 10 mM Tris-HCl pH 8.5) with 25 ml of solution per gram of pellet, then incubated at 25°C and 150 rpm for 10 min, and centrifuged at 4°C and 10000xg for 10 min. Immediately thereafter, the pellet was resuspended in osmotic solution 2 (10 mM MgSO_4_ in cold water) with 25 ml of solution per gram of pellet, then incubated at 4°C and 150 rpm for 10 min, and centrifuged at 4°C and 10000xg for 10 min. The periplasmic fraction was defined as the total activity found in the supernatants after centrifugation. The specific asparaginase productivity (U/g) was calculated as the ratio of catalytic activity (U) per biomass weight (g). Measurements were performed in triplicate. Cultures transformed with the pET-22b(+) plasmid were used as negative control.

### Purification

Purification was carried out by anion exchange chromatography [45,46], using a MonoQ 8-ml column attached to a AKTApurifier-UPC900 FPLC (GE Healthcare Bio-Sciences, USA). L-Asparaginase was extracted from the periplasmic space as detailed in the previous method section. The crude solution after osmotic shock was centrifuged and filtered through a 0.22 μm pore-size, then adjusted to pH 8.5 with Tri-HCl buffer. The MonoQ column was equilibrated with Tris-HCl buffer (50 mM pH 8.5), the crude L-asparaginase mixture was loaded into the column and washed with the same equilibration buffer. Elution was performed with a ramp of 0-100% 1 M NaCl in 50 ml. The presence of L-asparaginase was screened by measuring the catalytic activity.

### Asparaginase catalytic activity

Asparaginase catalytic activity was assayed by measuring the release of ammonia determined by direct nesslerization [47]. The enzymatic reaction was started by dispensing the substrate solution (10 mM L-Asn, 100 mM Tris-HCl pH 8.6) to the L-asparaginase samples up to 150 μl. All samples were assayed simultaneously in a 96-well-plate, using an Infinite M200PRO plate-reader with automatic dispensers (Tecan Trading AG, Switzerland). The samples and substrate solution were pre-heated at 37°C. The reaction was stopped after 10 min by dispensing 50 μl of TCA (0.3 M). The presence of ammonia was measured by the increase in absorption at 425 nm after the addition of 50 μl Nessler’s reagent. Ammonia concentration was determined with a calibration curve made with ammonium sulfate. Measurements were performed in triplicate.

### Total protein quantification

Total protein concentration was determined with Pierce^TM^ BCA Protein Assay Kit used as per manufacturer’s instructions and confirmed with the L-asparaginase absorption coefficient at 278 nm (E^1%^= 7.1) [48]. For samples that weren’t pure enough, SDS-PAGE gel densitometry analysis was used to the correct concentration values using the >99% pure bands as standard. Densitometry analysis was performed with the Gel Analyzer II software (Copyright license 2010) with rolling ball background subtraction.

### Gel electrophoresis

Native-PAGE and SDS-PAGE was performed using 4-20% polyacrylamide gels with the corresponding manufacturer’s running and loading buffers (Bio-Rad, USA), run at 150 V for 1 h. Samples of 5 μl were mixed at a 1:1 (v/v) ratio with the loading buffer and incubated for 5 min at 90oC in the case of denaturing SDS-PAGE. For Native-PAGE the samples were mixed with the native loading buffer and no further treatment was done. Gels were stained with 0.02% Coomassie Brilliant Blue G-250 and washed with de-staining solution (25 ml ethanol, 40 ml acetic acid, up to 500 ml with distilled water).

### Mass spectroscopy

Molecular weight of the native L-asparaginase and A38C-T263C mutant was obtained by mass spectroscopy. A MALDI 4800 plus TOF/TOF (SCIEX, USA) was used in positive linear mode, with sinapinic acid (10 mg/ml), in acetonitrile and 0.1% trifluoro acetic acid (50:50 v/v) as matrix. Calibration was performed with Cal Mix 3 (5735-66431 ±50 Da), then a commercial natural L-asparaginase II (*E. coli*) was set as standard (34600 ±50 Da).

Identity of the native L-asparaginase was confirmed using MS/MS analysis [49]. Briefly, the excised L-asparaginase band from SDS-PAGE was digested overnight at 4ºC with 50 μl of trypsin (13 ng/μl) and the peptides purified using a reverse phase C-18 zip-tip column. Eluted peptides were plated on the MALDI target plate mixed 1:1 (v/v) with α-cyano-4-hydrooxycinnamic acid (5 mg/ml) in acetonitrile and 0.1% trifluoro acetic acid (50:50 v/v) as matrix, and then analyzed in positive reflector MS/MS mode. Calibration was performed with Glu-Fib-B1 (1571.61 ± 0.5 Da). The precursor peptide pattern was compared against the *E. coli* taxonomy from the Mascot database online server.

### Site-specific PEGylation crosslinking

The site-specific PEGylation crosslinking was carried out by reacting the terminal maleimide groups of the PEG polymers with the surface-exposed cysteines on L-asparaginase using the well-known thiol-Michael addition click chemistry [36]. Briefly, L-asparaginase (A38C-T263C) in the crosslinking buffer (0.1 M potassium phosphate, 5 mM EDTA, 0.150 M NaCl, pH 6.5-7.0) was reduced with 5 mM TCEP for 30 min at room temperature. Immediately after, the reduced solution was desalted and the concentration of readily-exposed Cys was calculated using 5,5’-dithiobis-2-nitrobenzoic acid (Ellman’s reagent) [50]. The concentration of readily-exposed Cys was adjusted to 0.1 mM and reacted with the corresponding Bi-MAL-PEG polymer (1000, 2000 or 5000 g/mol) in a 40:1 molar ratio of PEG-to-Cys for 3 h at 25°C. Although a smaller molar ratio of PEG-to-Cys should promote greater likelihood of intramolecular crosslinking over intermolecular [51], preliminary experiments showed low recovery of neo-conjugates so we decided to keep a higher excess of PEG during the reaction. The reaction was stopped by addition of 10 mM DTT for 15 min, and the neo-conjugates were recovered by gel filtration chromatography or by repeated ultrafiltration steps in a 100 kDa cut-off filter. Since L-asparaginase subunits have a mass of ~35 kDa, non-conjugated subunits should pass through this membrane while tetrameric (or higher) conjugates are retained.

### Computational analysis

Geometry optimizations and electronic property computations of the model structure from L-asparaginase monomer were performed at the B3LYP/6-31G* level of theory using the Gaussian 09 program [52-54]. The model structure comprises the amino acids forming the natural disulfide bond and the serine mutations (C77-105S), along with four adjacent residues (D76, D78, K104 and D106). The starting atomic coordinates were extracted from the crystal structure reported by Swain *et al.* (1993) [1]. Partial optimizations, in which last carbon atoms in adjacent residues are fixed, were carried out to explore the effect of substituting the S atoms of cysteine in the natural disulfide bond by OH groups thus generating the serine residues. The solvent effect of water was considered by using the polarizable continuum model (PCM)/X throughout the computations [55].

## Results and discussion

### Characterization of site-specific PEGylation crosslinking

Site-specific PEGylation was performed using thiol-maleimide chemistry [36], to target cysteine residues previously introduced by mutagenesis at positions A38 and T263. These positions were selected in order to minimize the distance between Cys-to-Cys on adjacent subunits and maximize it within the same subunit. We argue that this promotes a higher ratio of intramolecular (within the L-asparaginase tetramer) as opposed to intermolecular (tetramer-to-tetramer) crosslinking. Selection of the mutation sites was done using the reported crystal structure coordinates from Swain *et al.* (1993). The distance between cysteine residues on adjacent subunits is 18-34 Å, while within the same subunit it is 55 Å. The selected cysteine mutations are far (~29 Å) from the active sites (**Fig 1A**) [1]. The tetrameric-structure of L-asparaginase has four active sites, each site is formed by the association of five residues from one subunit with two of the adjacent subunit (**Fig 1B**) [1,56]. It is difficult to select PEGylation positions further away from the active sites since L-asparaginase is a compact molecule. Nevertheless, the active site residues are relatively buried in comparison with the mutated cysteines (for PEGylation), the computed solvent accessible area of the catalytic center was 17.2 vs. 89.3 Å^2^ of the mutated positions (A38C-T293C), which reduces the likelihood of the active sites being affected after site-specific PEGylation.

**Fig 1.**
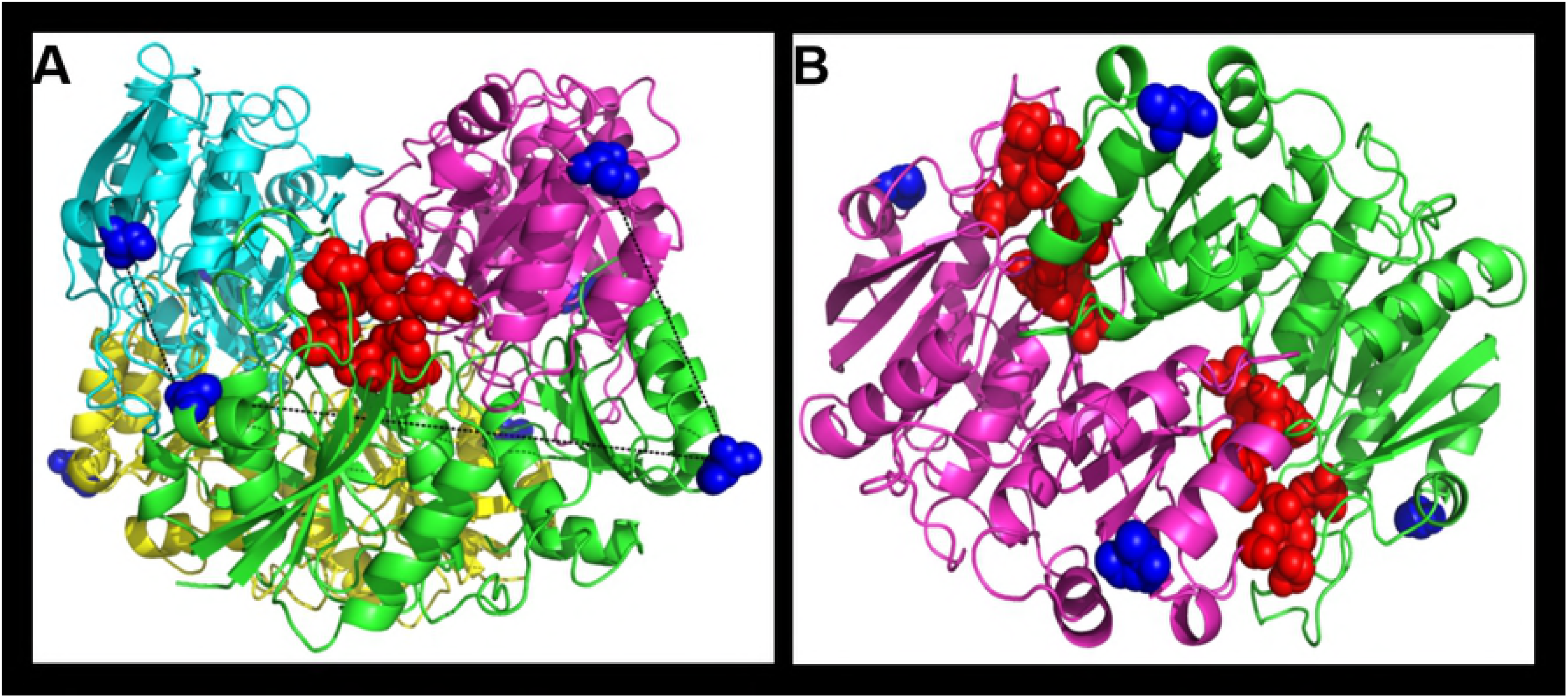
Selection of PEGylation sites on L-asparaginase. (A) L-Asparaginase tetramer with subunits highlighted with different colors, one active site is represented in red spheres, the pre-selected PEGylation positions in blue spheres, and the distances between mutated residues (A38C and T632C) are shown by black dotted lines. (B) L-Asparaginase dimer showing two active sites relatively buried (red spheres) in comparison with the PEGylations sites (blue spheres).

Site-specific PEGylation crosslinking was performed with homogeneous Bi-MAL-PEG polymers of 1000, 2000 and 5000 g/mol, which resulted in three physically different conjugates. Increasing the polymer size caused the neo-conjugate rigidity to decrease until it became completely soluble. The 1kDa-PEG-conjugate was a semi-solid gel (**Fig 2A**), while the 2kDa-PEG-conjugate was physically a soft gel in which a fraction of L-asparaginase (catalytic activity) remained in solution. The 5kDa-PEG-conjugate was a fully soluble construct. We think these results can be explained by the degree of dynamic freedom of the PEG molecule and kinetics of the thiol-maleimide reaction. The mutated cysteines on the L-asparaginase surface are the nucleophiles responsible for attacking the electrophilic carbon of the maleimide ring [36]. The first step of the crosslinking reaction proceeds fast in the presence of a Bi-MAL-PEG molar excess and at low protein concentration [51]. Once a cysteine reacts with one reactive side of the homo-bifunctional linker, the rate of the second reaction step depends on how fast the Bi-MAL-PEG finds the second Cys partner. For the small PEG polymer to react with the intramolecular partner is less probable than to react with an intermolecular thiol partner therefore the rate of intermolecular crosslinking is superior. The result is a hydrogel of intermolecularly crosslinked L-asparaginase. The hydrophilicity of PEGs also influences the rate of this second step [57], since interaction with water delays the finding of a second intramolecular partner. At increasing Bi-MAL-PEG length the neo-conjugates became more soluble because of the promotion of intramolecular over intermolecular crosslinking due to an increased dynamic freedom of PEG. Subunit crosslinking of L-asparaginase has been reported by Balcão *et al.* (2001), although no direct evidence of this crosslinking was shown [58]. Handschumacher and Gaumond (1972) also reported on L-asparaginase crosslinking that resulted mainly in dimers [59]. Our site-specific PEGylation strategy yielded a highly homogeneous and covalently crosslinked 2kDa-PEG-conjugate, which ran equal to the native L-asparaginase tetramer on a Native-PAGE (**Fig 2B**) and did not dissociate during denaturing SDS-PAGE electrophoresis (**Fig 2C**) proving that the subunits were indeed covalently crosslinked.

**Fig 2.**
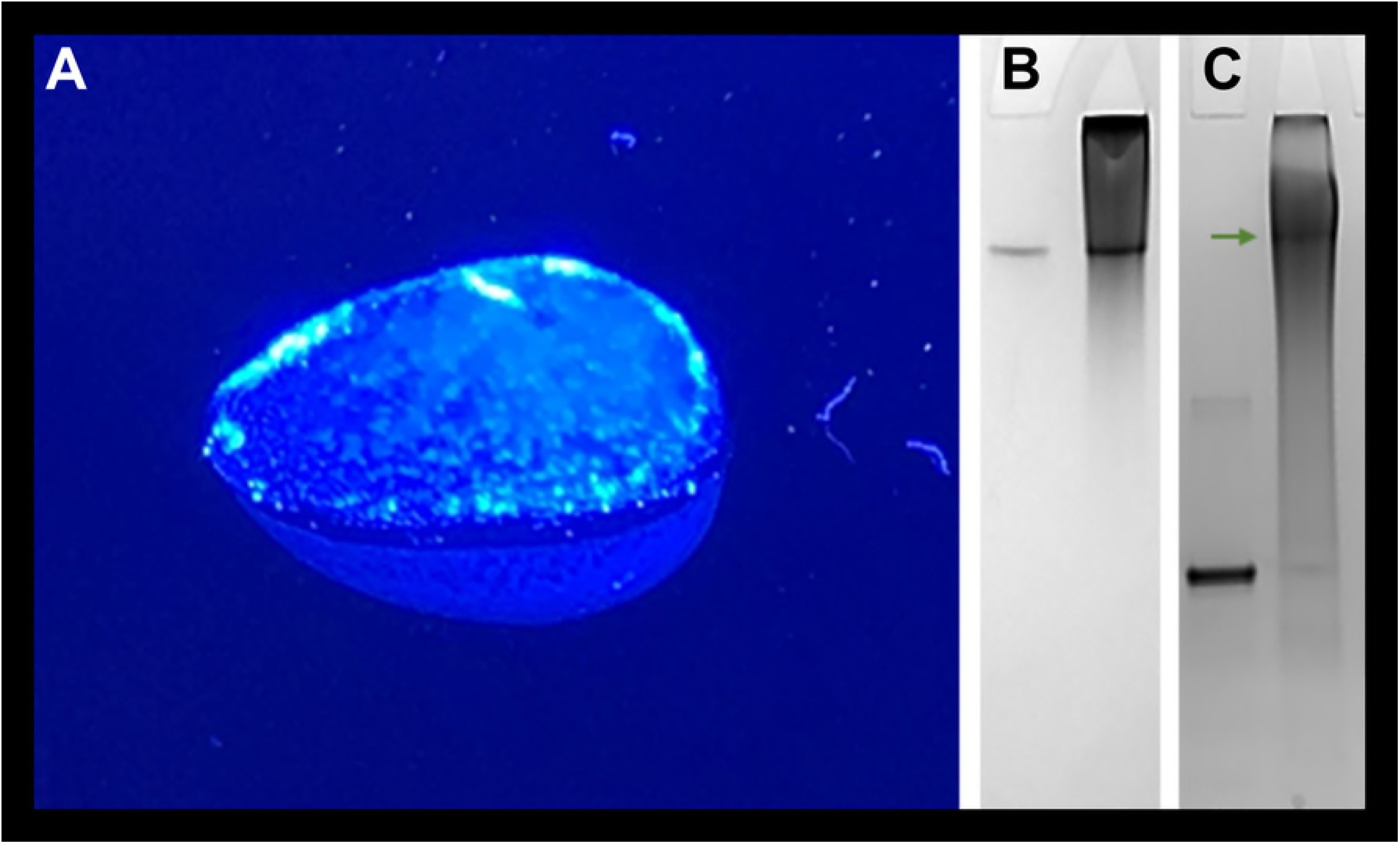
Site-specific PEGylation crosslinking with 1 and 2 kDa PEG. (A) 1kDa-PEG-conjugate gel. (B) Native and (C) SDS-PAGE of native L-asparaginase (left lanes) and the 2kDa-PEG-conjugate (right lanes). Note that a substantial amount of the 2kDa-PEG-conjugate did not enter the gel because it was too large.

Site-specific PEGylation with the 5000 g/mol PEG was characterized by size exclusion chromatography to qualitatively follow the covalent crosslinking of the subunits. The mutant L-asparaginase (A38C-T263C) was reduced with TCEP prior to and after PEGylation and then analyzed using gel filtration chromatography. Due to the quite low purity of the starting mutant solution, eluted fractions were also evaluated for catalytic activity to corroborate the presence L-asparaginase. Prior to PEGylation, it was observed that the mutant (A38C-T263C) eluted at several fractions in which more than 50% of the catalytic activity was found from 8-13 ml, while after PEGylation most of the activity was found at one fraction which eluted at 8-9 ml (**Fig 3**). The elution peak of the 5kDa-PEG-conjugated is approximately related to a relative molecular weight of 600-900 kDa, double of that reported for L-asparaginase randomly PEGylated using the lysine-directed approach [16,18]. One has to keep in mind that thiss result is not entirely accurate due to the high hydrophilicity of PEG that reduces protein mobility on gel filtration columns [57]. Despite of this, the calculated molecular weight was significantly higher than the 138.4 kDa of the native L-asparaginase tetramer. This implies that the 5kDa-PEG-conjugate likely also involves intra-intermolecular crosslinking. In any case, this neo-conjugate clearly exceeds the threshold of glomerular filtration which is beneficial for its application in leukemia [10,60], similar to the existing commercial randomly-PEGylated L-asparaginase [3-6,13]. The contrast between the catalytic activity eluted from gel filtration prior (wide distribution along 5 ml) and after (almost at a single 1 ml) PEGylation suggests that the 5kDa-PEG-conjugate must be indeed covalently crosslinked as it was not affected by reduction with TCEP contrary to the precursor mutant.

**Fig 3.**
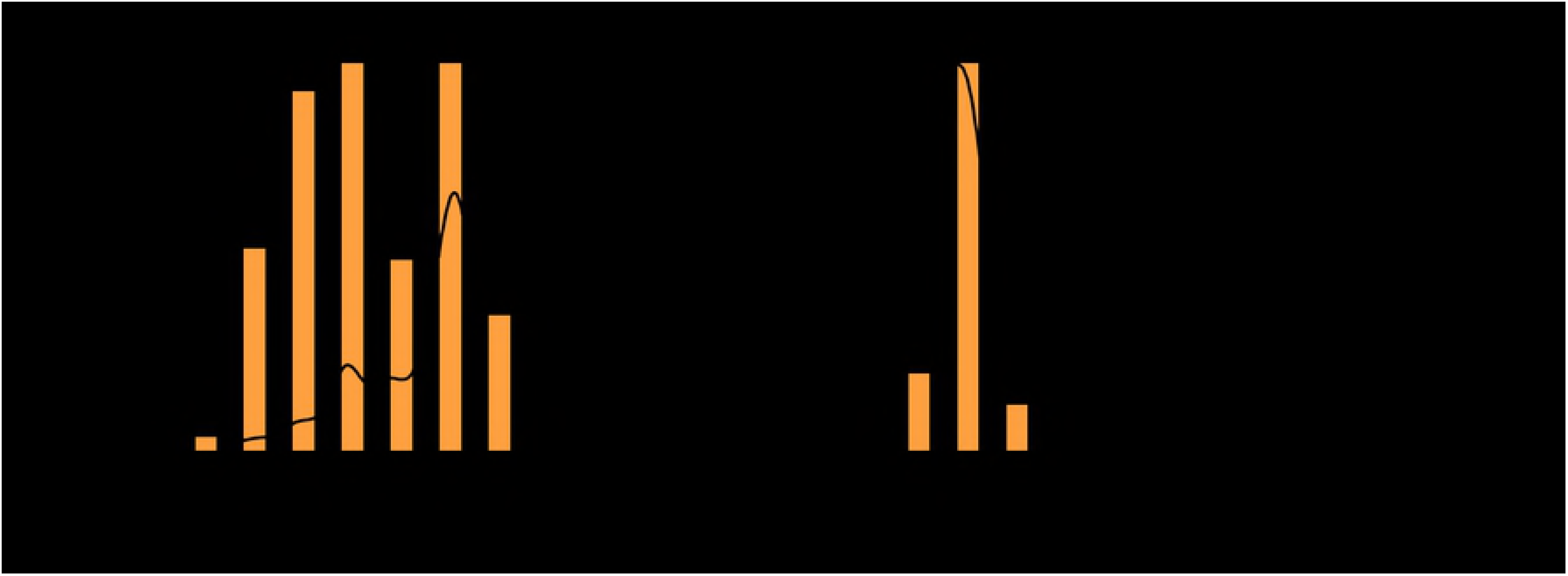
Site-specific PEGylation followed by size exclusion chromatography. (A) Mutant L-asparaginase (A38C-T263C) reduced with TCEP and eluted from the gel filtration column. Fractions 8-13 ml were pooled and used for the site-specific PEGylation reaction. (B) 5kDa-PEG-Conjugate after crosslinking reaction, reduced with TCEP and eluted from the gel filtration column. The continuous curve represents relative absorbance at 280 nm, the bars are the relative asparaginase catalytic activity of the collected fractions.

The product recovery after the site-specific crosslinking reaction by gel filtration was modest. The fraction where most of the 5kDa-PEG-conjugate eluted represents less than 20% of the initial catalytic activity exhibited by the starting mutant enzyme solution. For that reason, we also tested ultrafiltration for product purification using 10 and 100 kDa cut-off filters. The washing of the starting mutant enzyme solution after reduction with TCEP was done in a 10 kDa cut-off filter, immediately followed by the crosslinking reaction. The 5kDa-PEG-conjugate was pooled by performing repeated washing/concentration steps with a 100 kDa cut-off filter, since the non-conjugated L-asparaginase should flow-through these pores. The recovery by this method was about 48% of the initial catalytic activity **(S1 Table)**. This method is simpler and more efficient than the gel filtration approach **(S2 Fig)**.

Our results are in stark contrast with those reported for the established random Lys-directed PEGylation of the enzyme where multiple molecular weight conjugates of L-asparaginase are obtained thus complicating subsequent separation processes required for the commercialization of this drug. Such disadvantage of random PEGylation is exemplified in the report by Soares *et al.* (2002), where a wide distribution of molecular weight conjugates could be observed by SDS-PAGE electrophoresis [18]. In addition to the present article, the work by Balan *et al.* (2007) also highlights the specificity of Cys-directed PEGylation, although in this case the conjugation was performed at the natural disulfide bond [34].

### Effect of site-specific PEGylation crosslinking on catalytic activity

The antileukemia mechanism of L-asparaginase is still not fully understood, but there is a strong agreement that this enzyme acts by nutritional deprivation. Depletion of L-asparagine is perceived as the main trigger to destabilize the malignant cells, which are by themselves incapable of efficiently synthesizing this amino acid endogenously and therefore depend on the circulating L-Asn in blood. After administration of L-asparaginase, the enzyme reduces asparagine concentration in blood which impairs protein biosynthesis in the leukemia blasts and subsequently induces apoptosis in the cells that could not develop resistance [2-6]. Therefore, retention of the L-asparaginase catalytic activity after any modification is the most important *in vitro* parameter to extrapolate its potential antileukemia effect *in vivo*. PEGylation is the most utilized conjugation method to enhance pharmacokinetics of protein-based drugs [10], and among many examples, a commercial PEGylated L-asparaginase from *E. coli* is being used as frontline treatment [2,5]. However, PEGylation is also known for promoting restriction of protein structural dynamics thus diminishing the catalytic activity at an increasing degree of modification [14,15].

Because of its therapeutic importance L-asparaginase has been modified using several types of materials, e.g., bound to inorganic matrices, conjugated with natural and synthetic polymers, encapsulated into nanoparticles, and some hybrid-type approaches. A wide variety of results have been reported related to the pharmacodynamics parameters of those modifications, but in terms of PEGylation there is a clear conclusion that L-asparaginase catalytic activity is affected when conjugated using a Lys-directed approach depending on the degree of modification [61]. Wang *et al.* (2012) demonstrated that Lys-directed PEGylation can be improved through an alkylation strategy, the PEGylated L-asparaginase was highly homogeneous but still the catalytic activity was reduced to 44% compared to the non-modified enzyme [19]. Alkylation allows to control the degree of modification tightly, but PEGylation sites could not be selected which might explains the drastic reduction in catalytic activity even with a low degree of modification. Balan *et al.* (2007) successfully modified L-asparaginase at a single site using a three-carbon bridge strategy. The PEG neo-conjugate retained full catalytic activity independently of the PEG molecular weight. Unfortunately, this strategy is limited by the relative solvent accessible area of the natural disulfide bonds in the protein. The PEGylation sites cannot be selected which limits the potential to hide proteolytic and antigenic epitopes [34]. Another important aspect is that L-asparaginase is a tetrameric enzyme and the association of the monomeric subunits is imperative to form the active sites and exhibit catalytic activity [1,56]. It has been shown that PEGylation reduces the ability of this enzyme to form its tetrameric-structure [18], which could reduce the efficiency of this drug since the average administration is around 7 μg of L-asparaginase per milliliter of blood [5,62]. A covalent crosslinking strategy of L-asparaginase subunits has been proposed to stabilize the active multimeric structure, but previous results showed that this approach is linked to a cost in catalytic activity due to the Lys-directed conjugation employed. Balcão *et al.* (2001) retained 35% activity [58], while Handschumacher and Gaumond (1972) only managed to keep 17% compared to the non-modified L-asparaginase [59].

In this work we performed site-specific PEGylation of L-asparaginase subunits that yielded a fully soluble neo-conjugate after crosslinking with 5000 g/mol Bi-MAL-PEG. The average catalytic activity of the 5kDa-PEG-conjugate and its non-conjugated mutant precursor (A38C-T263C) was assayed *in vitro* by measuring the rate of L-Asn hydrolysis. L-asparaginase concentration was selected to be 14 μg/ml to simulate a therapeutically relevant concentration. The precursor mutant exhibited lower catalytic activity than the native non-modified L-asparaginase (P<0.05), 116 ± 6 vs. 161 ± 9 U/mg, respectively (**Table 1**). This was unexpected since the mutated cysteines are relatively far away from the active site (**Fig 1**), which suggest that the high solvent accessible area of the mutated residues (89.3 Å^2^) might promote disulfide subunit multimerization that eventually affects formation of the L-asparaginase tetrameric-structure required to perform catalysis [18,34,63,64]. After site-specific crosslinking, the 5kDa-PEG-conjugate showed superior catalytic activity than the native L-asparaginase (P<0.05), 210 ± 11 vs. 161 ± 9 U/mg, respectively (**Table 1**). It was expected that this mutant should retain the same catalytic activity after site-specific PEGylation [34], but not that it would exceed the activity of native L-asparaginase. Since no mutations were performed near the active sites, these results support the hypothesis that the tetrameric-structure formation of L-asparaginase is a determinant step for its catalysis [18,34,63,64]. We presume that the covalent crosslinking maintains the subunits within high proximity between each other, hereby facilitating the formation of the active tetrameric-structure triggered by the presence of the L-Asn substrate [63,64], a process that usually is rate limiting. To support these observations we compared the catalytic activity of the 5kDa-PEG-conjugate and native non-modified enzyme at below biological temperature, to introduce an energetic effect that should delay the association of the tetrameric-structure thus reducing the average catalytic activity as well [17,40,64]. At 24°C the native L-asparaginase exhibited a catalysis of 47.2 ± 1.7 U/mg that is 30% of the one observed at 37°C. The activity obtained for the 5kDa-PEG-conjugate was 210 ± 6 U/mg, basically the same as observed at 37°C. This strongly suggest that the covalent intrasubunit crosslinking is playing a stabilazing effect that accelerates the formation of the active structure upon addition of the L-Asn substrate [63]. It is important to explain that due to the low purity of the starting mutant solution (A38C-T263C), the 5kDa-PEG-conjugate solution is not pure as well **(S2 Fig)**. This means that the L-asparaginase concentration value used is overestimated, which causes the value of the calculated specific activity to be underestimated **(S3 and S4 Tables)**.

**Table 1.**
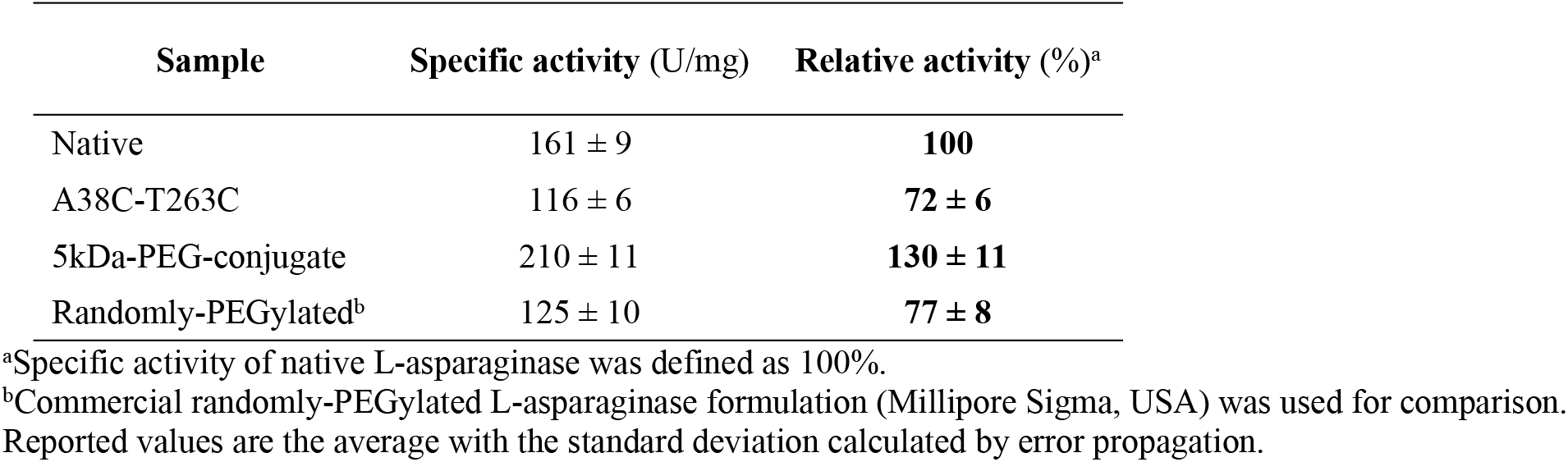
Catalytic activity of site-specific crosslinked L-asparaginase.

| Sample | Specific activity (U/mg) | Relative activity (%)<sup>a</sup> |
| --- | --- | --- |
| Native | 161 ± 9 | **100** |
| A38C-T263C | 116 ± 6 | **72 ± 6** |
| 5kDa-PEG-conjugate | 210 ± 11 | **130 ± 11** |
| Randomly-PEGylated<sup>b</sup> | 125 ± 10 | **77 ± 8** |

<sup>a</sup>Specific activity of native L-asparaginase was defined as 100%.
<sup>b</sup>Commercial randomly-PEGylated L-asparaginase formulation (Millipore Sigma, USA) was used for comparison.
Reported values are the average with the standard deviation calculated by error propagation.

In summary, a new site-specific PEGylation strategy to crosslink L-asparaginase subunits generated higher molecular weight neo-conjugates and improved the average catalytic activity likely involving an intrasubunit proximity-stabilization mechanism. Compared to previous reported PEGylation strategies, our PEGylation method is the only one that improved the catalytic potential of L-asparaginase (**Table 2**). A similar method has not been proposed before possibly due to the difficulty to express and purify recombinant L-asparaginase containing non-natural cysteines [42]. We are the first to report this kind of mutation in L-asparaginase designed for site-specific conjugation, which is beneficial to improve this therapeutic enzyme since a commercial recombinant L-asparaginase is already on the market [2].

**Table 2.**
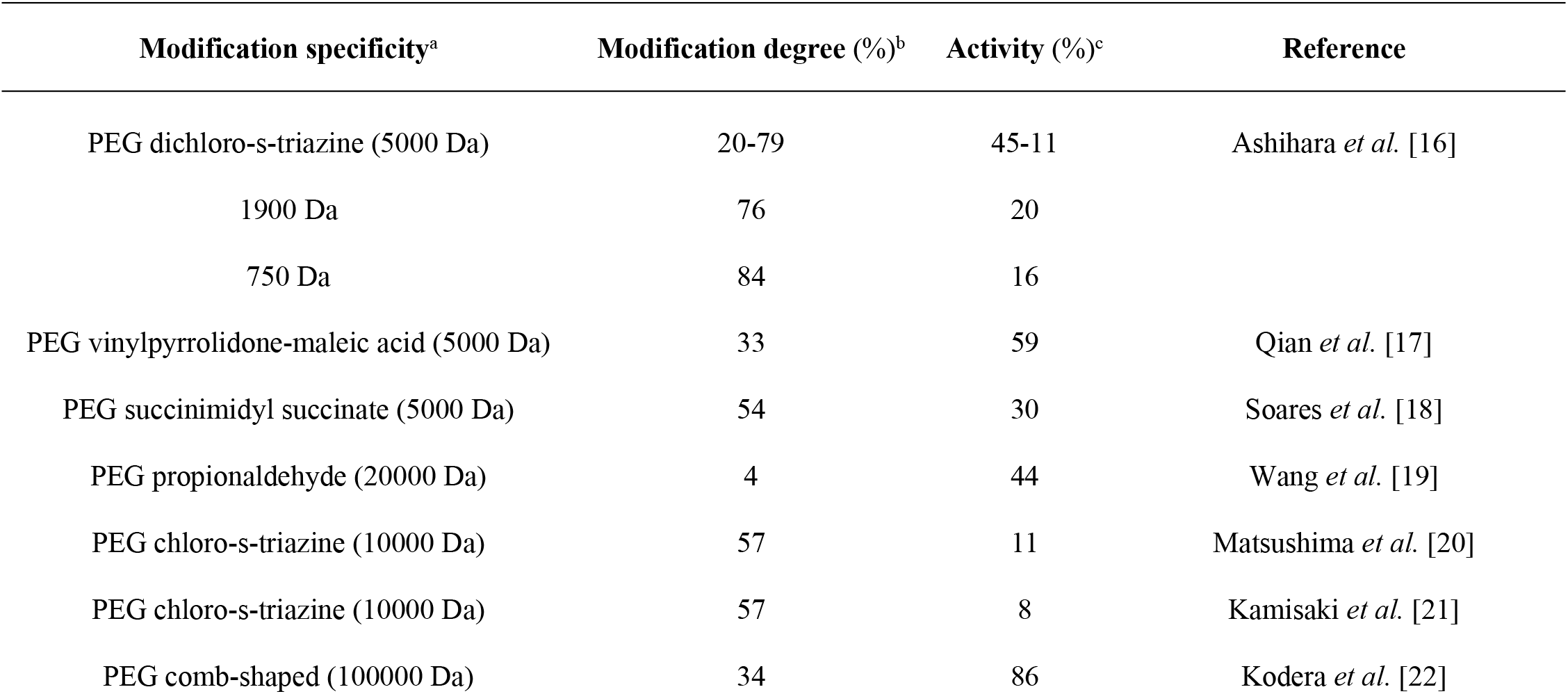
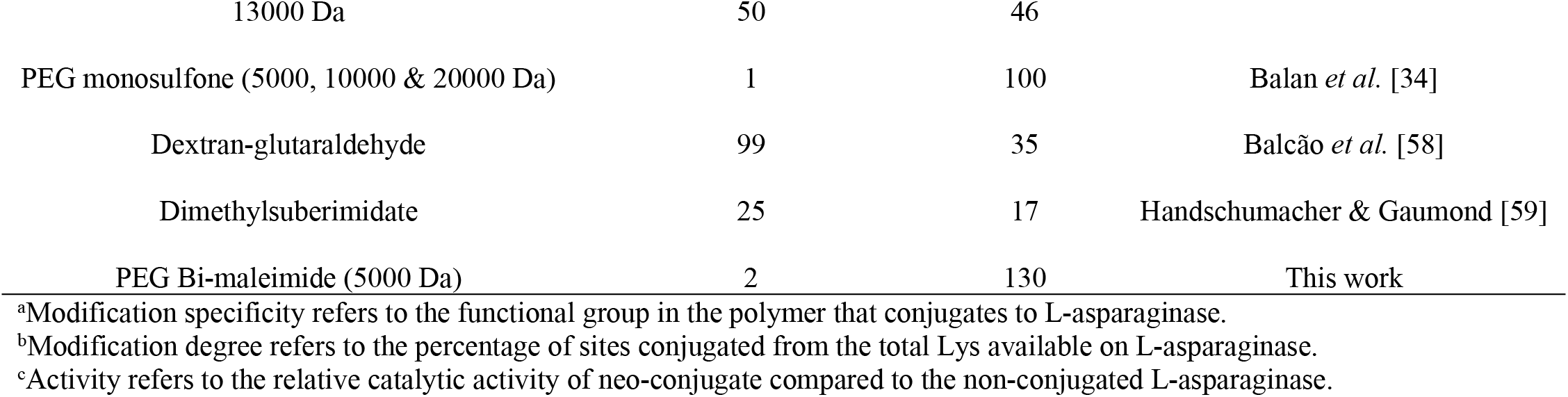
Comparison of PEGylation methods for L-asparaginase.

Studies on PEGylation of L-asparaginase show a fine trend of decreasing catalytic activity with increasing degree of modification (**Fig 4**), which can be explained by the restriction of protein structural dynamics related to the number of polymers conjugated to the protein, but independently of the PEG molecular weight [14,15]. Our results do not deviate from this trend, but we rather report a flexible strategy that allows to vary the conjugation site to target potential proteolytic and immunogenic epitopes. Simultaneously, the covalent subunits crosslinking increases the molecular weight to delay glomerular filtration [10,60].

**Fig 4.**
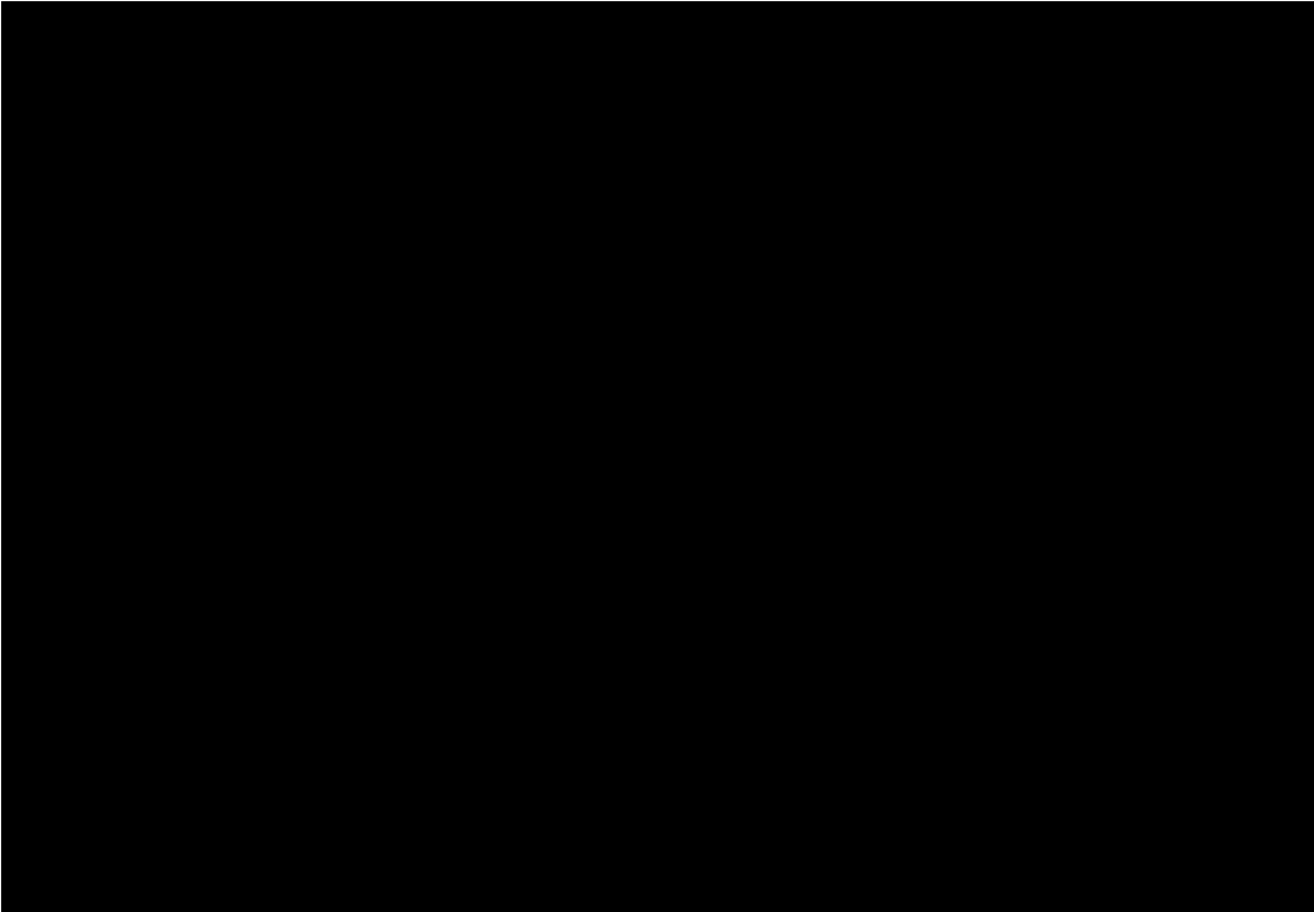
Trend on catalytic activity with degree of PEGylation. Reported catalytic activity of PEGylated L-asparaginase with degree of conjugation. Relative activity is expressed as percentage compared to the non-conjugated L-asparaginase. Modification degree refers to the percentage relative to the available lysines. The results to generate this figure were extracted from references [16-22,34] and this work.

L-Asparaginase is also being studied as a possible solution to reduce the formation of acrylamide in fried food products [7]. Even though it is not the main scope of this report, the physical properties of our 1kDa-PEG-conjugate could be beneficial for industrial applications that use L-asparaginase. After treating raw products with L-asparaginase to hydrolyze the acrylamide precursor asparagine, the enzyme is difficult to recover due to its soluble state. Our neo-conjugate is a semi-solid gel that can be recovered from solution and reutilized. We tested the catalytic activity of this neo-conjugate within a temperature range from 20°C to 80°C to evaluate its potential for being used in industrial applications. The same 1kDa-PEG-conjugate sample was used throughout the whole experiment by washing the gel with distilled water after each catalytic cycle. Our neo-conjugate proved to be highly reusable along the 27 cycles tested, with a maximum catalytic temperature up to 60°C (**Fig 5**). Although L-asparaginase is a multimeric protein with the catalytic core in the middle of two subunits [1,56], our discreet conjugation didn’t affect its enzymatic function even in an insoluble state. This conjugation strategy could be extended to other industrially important enzymes to improve cost-efficiency.

**Fig 5.**
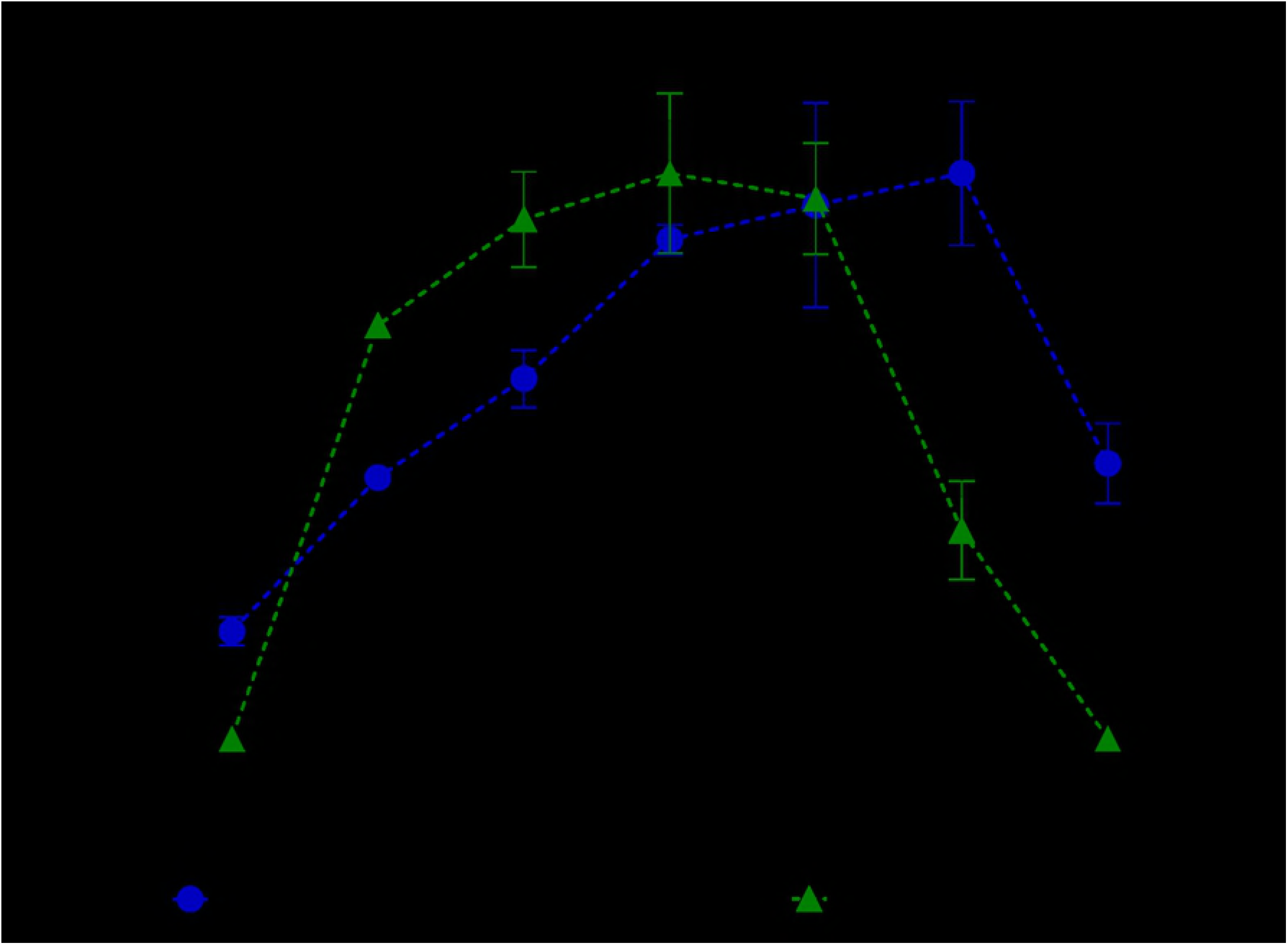
Reusability of the 1kDa-PEG-conjugate. Relative catalytic activity at temperatures ranging from 20°C to 80°C. The 1kDa-PEG-conjugate (in green), native L-asparaginase (in blue), and a commercial randomly-PEGylated L-asparaginase formulation (Millipore Sigma, USA) (in black). Relative catalytic activity was calculated by the absorption at 425 nm divided by the maximum signal for each sample and expressed as percentile (%). Reported values are average with error bars representing the 95% confidence interval. For the 1kDa-PEG-conjugate the same sample (semi-solid gel) was used throughout all the experiment.

### Secretory expression

In this work, a recombinant L-asparaginase was expressed using the commercial pET system (Novagen, Germany). Induction of the pET22b-AspII expression system with IPTG concentrations of 500 and 1000 μM was counterproductive, since it significantly decreased the secretion of native L-asparaginase in comparison with induction using 10 and 100 μM (P <0.01) **(S5 Fig)**. These results agree with other studies using the same pET system. Khushoo *et al.* (2004) found that a concentration of 100 μM of IPTG was favorable for secretion and cell growth [37], while Vidya *et al.* (2011) reported the highest level of secretion inducing with 10 μM of IPTG [39]. Extracellular secretion of L-asparaginase to the growth medium was monitored at 8, 16 and 24 h post-induction. Most of the catalytic activity was found in the culture medium at 8 h post-induction, afterwards the ratio of activity-to-biomass decreased **(S5 Fig)**. It is unnecessary to grow cultures beyond 8 h, which is a practical information when producing high amount of this enzyme recombinantly. Due to the convenience of producing L-asparaginase as secretory product to avoid formation of inclusion bodies [37-41], we tested the ratio of extracytoplasmic secretion to the periplasmic space and culture medium for three recombinant constructs: the native L-asparaginase, the double mutant A38C-T263C, and a C77-105S mutant that lacks the natural disulfide bridge. The presence of catalytic activity was measured in the clear culture medium after biomass centrifugation and in the periplasmic space fraction obtained by osmotic shock. It was found that for each independent construct the secretion ratio did not vary significantly, a similar amount of catalytic activity was found in both the culture medium and periplasmic fraction. On the contrary, the absolute productivity (U/g) varied between the three constructs. The native construct exhibited ~4.1-fold secretory expression compared to the A38C-T263C mutant, while the C77-105S mutant showed almost no expression (**Table 3**).

**Table 3.** Extracytoplasmic secretion of recombinant L-asparaginase constructs.

| Sample | Culture medium |  | Periplasmic space |  |
| --- | --- | --- | --- | --- |
|  | Productivity (U/g) | Ratio (%)<sup>b</sup> | Productivity (U/g) | Ratio (%)<sup>b</sup> |
| Native | 1273 ± 74 | 40.3 ± 6.4 | 1883 ± 152 | 59.7 ± 6.3 |
| A38C-T263C | 324 ± 35 | 42.9 ± 9.8 | 431 ± 20 | 57.1 ± 5.3 |
| C77-105S<sup>a</sup> | 31 ± 7 |  | 13 ± 2 |  |

<sup>a</sup>The C77-105S mutant exhibited insignificant signal and thus the secretion ratio was not calculated.
<sup>b</sup>Secretion ratio was calculated as percentage of total extracytoplasmic secretion.
Reported values are the average with standard deviation.

The results from secretory expression suggest that the presence of cysteine residues in L-asparaginase is a factor inhibiting secretion of the enzyme, despite of the constructs all having the natural signaling peptide [43]. The almost-full inhibition of extracytoplasmic secretion (~70 times less than the native) by the C77-105S mutation may be related to the mechanism used by *E. coli* to translocate the recombinant protein through the cytoplasmic membrane. Introduction of additional cysteines (A38C-T263C mutant) while still maintaining the natural disulfide bridge (between C77 and C105) affected secretion to a lesser degree, which may be just an inherent consequence of a lower Cys-tRNA abundance in the expression organism [42]. The question that now arises is whether the position of the natural cysteines plays any important role in L-asparaginase secretion, or if solely the presence is sufficient. This is the first study that reports removal or introduction of cysteine residues in L-asparaginase and its effect on extracytoplasmic expression.

### Effect of natural disulfide bond mutation on catalytic activity

The specific catalytic activity of the recombinant L-asparaginase C77-105S mutant was determined to evaluate the feasibility of using this mutant in future studies. It is possible to conjugate the natural cysteines in L-asparaginase (C77 and C105) as reported by Balan *et al.* (2007) [34], so these residues could interfere with the thiol-maleimide reaction targeted at canonical cysteines introduced by mutagenesis. For this reason, it is beneficial to understand the importance of this natural disulfide bridge on the catalytic activity of L-asparaginase. This C77-C105 bond is relatively away from the active site (~35 Å) [1], and does not significantly impact the catalytic activity after been reduced [65]. The solvent accessible area decreases upon formation of the tetrameric-structure suggesting that hydrophobic interactions on the internal subunits interface stabilize the quaternary structure [64], while binding of the L-Asn substrate promotes additional structural stabilization [63]. Therefore, elimination of the natural disulfide bond should not significantly affect the catalytic activity. We replaced the cysteine amino acids (C77 and C105) by two serine residues with the intention of minimizing any alteration of the surrounding environment of these residues. We found no significant difference in the specific activity exhibited by the native L-asparaginase and C77-105S mutant (**Table 4**). This implies that removal of the natural disulfide bond should not destabilize the tetrameric-structure of L-asparaginase.

**Table 4.** Catalytic activity of L-asparaginase C77-105S mutant.

| Sample | Specific activity (U/mg) | Relative activity (%)<sup>a</sup> |
| --- | --- | --- |
| Natural | 177 ± 11 | **100** |
| Native | 156 ± 14 | **88 ± 10** |
| C77-105S | 160 ± 9 | **91 ± 7** |

<sup>a</sup>Specific activity of commercial natural L-asparaginase II (Millipore Sigma, USA) was defined as 100%. Reported values are the average with the standard deviation calculated by error propagation.

Computational analysis was used to evaluate possible structural disturbances caused by the C77-105S mutation in the vicinity of the natural disulfide bridge. The original structure from Swain *et al.* (1993) was used as starting coordinates for the natural cysteines (C77 and C105) along with four residues (D76, D78, K104, and D106) contained in the model [1]. After this model was geometrically optimized, the sulfur atoms in the cysteines were replaced by hydroxyl groups to generate the serine residues, and the resulting structure was optimized. Interestingly, the C77-105S mutation does not cause significant structural disturbance, as revealed by the superposition of the two optimized structures (**Fig 6**). It is highly possible that one hydrogen bond (1.828 Å O…HO) is formed between the serine residues. More details are presented in the supporting information **(S6 Fig and S7 Table)**. These calculations agree with the catalytic activity observed for the C77-105S mutant, showing that no structural disturbance takes place hence the active site should not be affected. These findings support the hypothesis that formation of the tetrameric-structure of L-asparaginase should be driven by other forces such as hydrophobic interactions and substrate biding [18,63,64].

**Fig 6.**
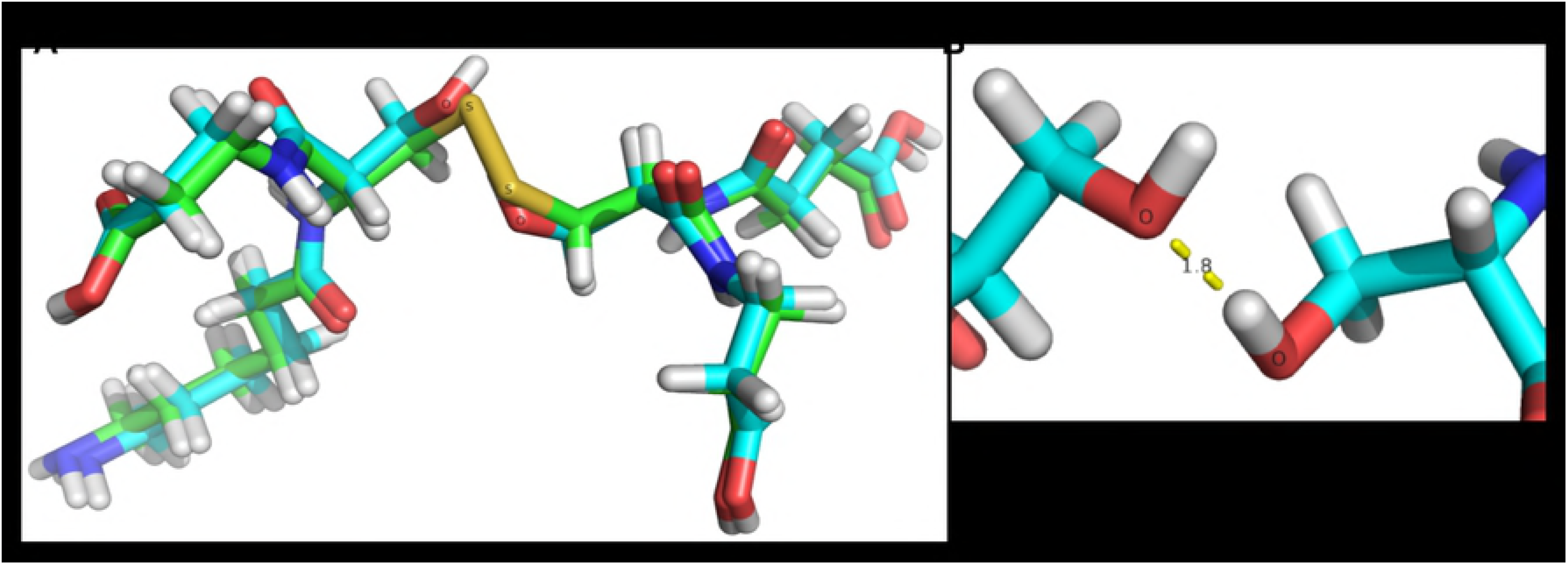
Computational analysis of the C77-105S mutation. (A) Superposition of geometrically optimized structures of the natural disulfide bond and C77-105S mutation. (B) Formation of a new hydrogen bond in the C77-105S mutant.

### Purification

The purification of recombinant L-asparaginase constructs was performed by a single step of anion exchange chromatography [45,46], using a high resolution MonoQ 8-ml column. The pellet from a 200 ml-culture sample was subjected to osmotic shock, and the clear crude mixture was purified on the MonoQ colunm. The native L-asparaginase was successfully purified with >95% purity (by SDS-PAGE) along three 1 ml-fractions **(S8 Fig)**. The final overall yield for this construct was 29 mg of pure L-asparaginase per liter of culture, which is an excellent although not as good as in previous reports using affinity purification methods [37,38,40]. Unfortunately, a His-tag affinity purification has been reported to interfere with the catalytic activity of L-asparaginase [38] and for this reason we designed our expression system with the intention of producing a native L-asparaginase without any modification. The mutant A38C-T263C was also purified with a good recovery although the purity was inferior (65% by SDS-PAGE) compared to native L-asparaginase **(S9 Fig)**. This was unexpected since both constructs have the same calculated isoelectric point (pI 5.8). At this point, it can only be assumed that the non-natural cysteines in the mutant may interfere in some way with the column binding. By last, the mutant C77-105S was purified with a very low yield, although this was expected because of the poor secretory expression observed (**Table 3**), the purity was also inferior compared to the other constructs (35% by SDS-PAGE) **(S9 Fig)**.

The relative molecular weight of the recombinant L-asparaginase constructs was calculated by mass spectroscopy. The main *m*/*z* peak was observed at 34605 g/mol for the native L-asparaginase, while 34634 g/mol for the mutant A38C-T263C (**Fig 7**). Unfortunately, it was not possible to obtain detailed information for the mutant C77-105S due to the limited amount of sample and the level of impurities. These results agree with the molecular weight calculated theoretically for the native L-asparaginase and mutant A38C-T263C, i.e., 34.60 and 34.63 kDa respectively. Both constructs were purified in its mature form since the pre-L-asparaginase should have a subunit mass around 36.8 kDa [43], we concluded that the natural signaling peptide was properly excised.

**Fig 7.**
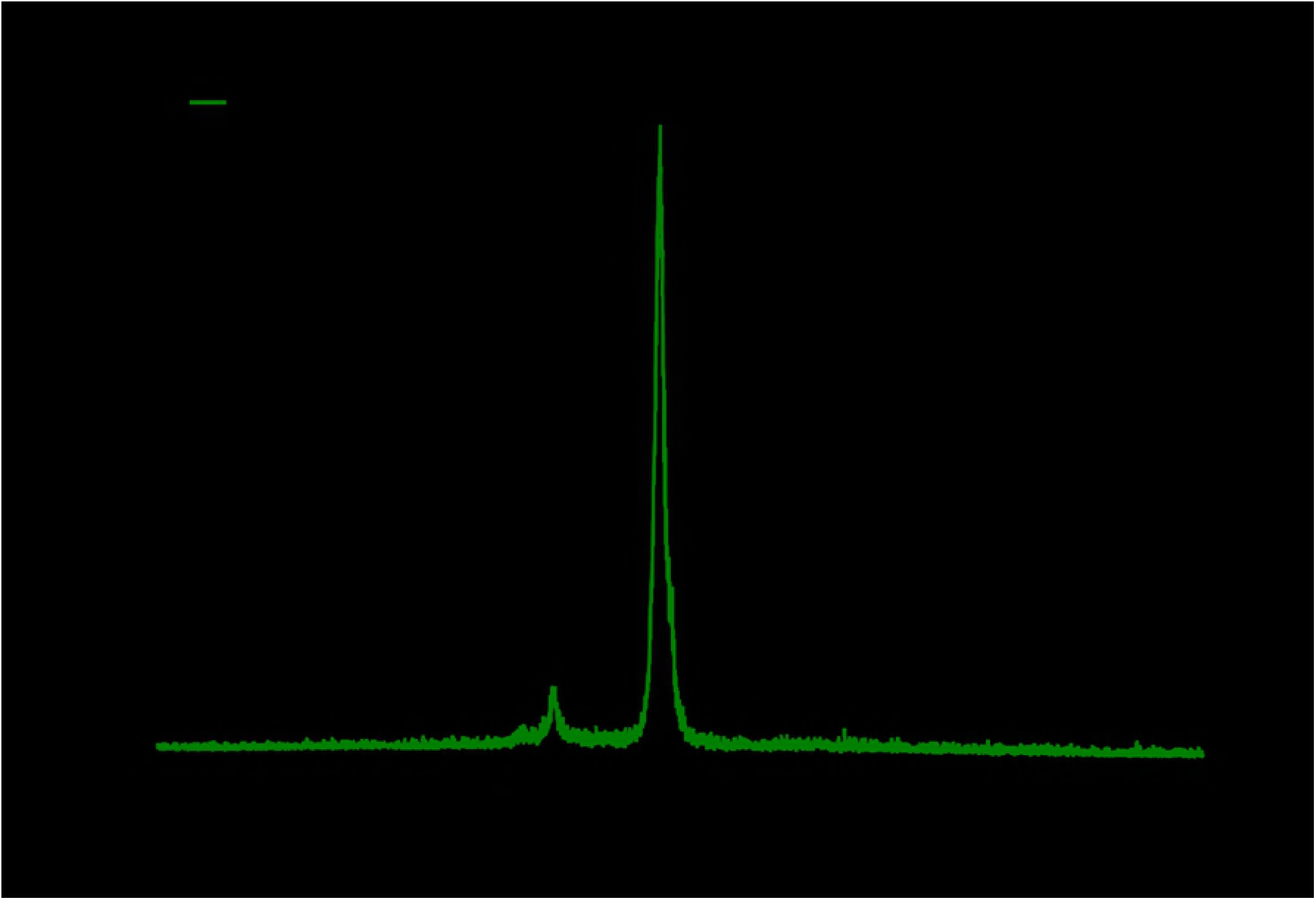
Subunit mass of recombinant native L-asparaginase and mutant A38C-T263C. The *m*/*z* peak for the native L-asparaginase subunit (in green) was observed at 34605 g/mol, while the *m*/*z* peak for the mutant A38C-T263C subunit (in black) was observed at 34634 g/mol.

MS/MS analysis was performed to the recombinant native L-asparaginase to confirm its identity. The peptides *m*/*z* pattern matched the theoretical trypsin-digested precursors pattern of *E. coli* L-asparaginase II from the Mascot database with a protein-score of 118 and 46% sequence coverage **(S10 Fig)**. In the Mascot database, protein-scores are calculated as −10·log(R), with R being the probability that the observed pattern is a random match. A score greater than 74 are considered significant (P<0.05).

## Conclusions

Herein we report an alternative site-specific conjugation strategy to modify the therapeutic enzyme L-asparaginase. Our strategy is based on three main pillars: firstly, that the molecular weight of L-asparaginase can be increased by crosslinking the subunits without the need to modify this protein excessively; secondly, that the polymers can be linked to the enzyme at pre-selected non-natural residues along the surface, to hide potential proteolytic and immunogenic epitopes; and thirdly, that intramolecular crosslinking of the subunits stabilizes the active tetrameric-structure to enhance catalytic activity. We demonstrated that site-specific PEGylation can be used to conjugate L-asparaginase at two different positions on its surface, while at the same time maintaining its full enzymatic potential. We found that introduction or removal of cysteine residues seems important for the proper extracellular expression of L-asparaginase in *E. coli* and achieved to purify this recombinant protein using a single anion exchange step.

Our 5kDa-PEG-conjugate was fully soluble with a molecular weight about double of randomly-PEGylated formulations, which is beneficial to delay glomerular filtration during the *in vivo* therapeutic application of this enzyme. Even more important, this neo-conjugate retained full catalytic activity, and we proposed that this is possible due to a tetrameric-structure stabilization effect that accelerates the formation of the active quaternary structure of L-asparaginase. In addition, the 1kDa-PEG-conjugate was a highly reusable semi-solid gel that exhibited catalytic activity and can be used for applications in the food industry. The reported site-specific conjugation approach provides the possibility to modify L-asparaginase pharmacokinetics while maintaining its pharmacodynamics properties, which is beneficial for the application of this drug in biological conditions.

## Acknowledgments

This work was developed with materials, instrumentation and facilities provided by Kai Griebenow’s research laboratory at Facundo Bueso building at the University of Puerto Rico at Rio Piedras campus, the Molecular Sciences Research Center at the UPR, and RISE Program at UPR Rio Piedras NIH Grant No. 5R25GM061151-16. We like to thank Arthur D. Tinoco for his help with the mass spectroscopy instrumentation.

## Supporting information

S1 Table. Recovery of site-specific PEGylation performed by ultrafiltration.

S2 Fig. Densitometry analysis of SDS-PAGE gels.

S3 Table. Correction of L-asparaginase concentration derived from densitometry analysis.

S4 Table. Calculation of asparaginase specific catalytic activity.

S5 Fig. Optimization of secretory expression.

S6 Fig. Geometrically optimized model structures for C77-105S mutation.

S7 Table. Geometrically optimized bond lengths and angles of model structures.

S8 Fig. Purification of recombinant native L-asparaginase.

S9 Fig. Purification of L-asparaginase mutants.

S10 Fig. Precursor peptides pattern of the recombinant native L-asparaginase.

